# Socio-cultural practices may have affected sexual dimorphism in stature in Early Neolithic Europe

**DOI:** 10.1101/2023.02.21.529406

**Authors:** Samantha L Cox, Nicole Nicklisch, Michael Francken, Joachim Wahl, Harald Meller, Wolfgang Haak, Kurt W Alt, Eva Rosenstock, Iain Mathieson

**Affiliations:** Department of Genetics, Perelman School of Medicine, University of Pennsylvania, Philadelphia, PA, USA; Physical Anthropology Section, Penn Museum, University of Pennsylvania, Philadelphia, PA, USA; Center of Natural and Cultural Human History, Danube Private University, Krems-Stein, Austria; State Office for Cultural Heritage Management Baden-Württemberg, Osteology, Konstanz, Germany; Institut für Naturwissenschaftliche Archäologie, AG Paläoanthropologie, Tübingen, Germany; State Office for Heritage Management and Archaeology Saxony-Anhalt, State Museum of Prehistory, Halle, Germany; Department of Archaeogenetics, Max Planck Institute for the Science of Human History, Jena, Germany; Bonn Center for ArchaeoSciences, Universität Bonn, Bonn, Germany

## Abstract

The rules and structure of human culture impact health and disease as much as genetics or the natural environment. To study the origin and evolution of these patterns, we take a multidisciplinary approach combining ancient DNA, skeletal metrics, paleopathology, and stable isotopes. Our analysis focuses on cultural, environmental, and genetic contributions to variation in stature in four populations of Early Neolithic Europe. In Central Europe, low female stature is likely due to male preference in resource allocation under conditions of stress. In contrast, shorter male stature in Mediterranean populations may reflect a lack of preference. Our analysis suggests that biological consequences of sex-specific inequities can be linked to culture as early as 7000 years before present. Understanding these patterns is key to interpreting the evolution of genetic and socio-cultural determinants of health, and our results show that culture, more than environment or genetics, drove height disparities in Early Neolithic Europe.

## 1 Introduction

Human skeletal variation reflects varying combinations of genetic, cultural, and environmental factors. While there are many links between culture and health in the modern world, the history and evolution of these relationships are not always well established. Due to the entanglement of these factors, our ability to draw conclusions about their effects has been limited in archaeological data. With the recent advent of ancient DNA sequencing technology, genetic information from ancient populations has become increasingly available. However, attempting to analyze changing patterns of variation based solely on genetic data is difficult—genotypes do not necessarily equate to phenotypes due to the effect of the environment. ^1^ Similarly, while it is tempting to predict phenotypic changes in ancient people based on their genetic variation, this is currently challenging as genetic effects are not always transferable across populations. ^2^ Our solution is to integrate these complementary fields to construct multidisciplinary analyses with phenotype, genotype, culture, and environment data from ancient human populations. This approach allows us to begin to separate the effects of these variables and reveal the interactions between genes, environment, and culture which are critical in shaping human health and variation.

Many traits of interest, including height, are highly polygenic, with thousands of independent genetic variants contributing significantly to heritability. One common approach to addressing the role of genetics in morphological change is to compare patterns of phenotypic variation with genetic ancestry or genome-wide patterns of genetic variation. ^3–5^ However, even for highly polygenic traits like height, genome-wide variation may not be directly relevant, leading to spurious associations between genetic effects, ancestry, and environmental confounds. For example, if a population is tall and has a high proportion of ancestry from Neolithic sources, it could be concluded that Neolithic ancestry is associated with “genetic tallness”; however, the effects could equally be non-genetic and related to lifestyle changes associated with agriculture. An alternative approach is to focus only on genetic variation that is known to be associated with a specific trait. ^6, 7^ Effect sizes for these trait-related variants estimated from genome-wide association studies (GWAS) of present-day individuals can be combined with genetic data from ancient individuals to calculate polygenic risk scores (PRS), which can be thought of as estimated genetic values for the phenotype. In European ancestry populations, polygenic scores for height can explain up to 25% of phenotypic variation in present-day individuals, ^8^ and 6-8% of variation in ancient individuals. ^9, 10^ On a broad scale, temporal changes in polygenic score over time in Europe are qualitatively consistent with changes in stature as inferred from the skeletal record, ^11^ while local deviations from this pattern provide evidence of environmental effects. ^10, 11^

Analyses of human populations over tens of thousands of years involve individuals that are diverse in genetic ancestry, environment, and culture and it is challenging to exclude the possibility of confounding by unmeasured variables. We therefore focus specifically on the European Early Neolithic. One of the most studied periods in prehistory, it represents a fundamental shift in technology, culture, and genetics. In particular, the *Linearbandkeramik* (LBK) culture of Central Europe is one of the most comprehensively documented Early Neolithic cultures, with an abundance of excavated settlements and cemeteries. ^12^ LBK groups tended to choose settlement locations based on the presence of rich loess soils for farming, and the northern edge of these soils appears to delineate the northern limit of LBK sites.^13, 14^ Bioarchaeological evidence indicates broad regional differences between individuals from northern settlements in this agricultural boundary zone vs southern settlements in a climate zone that was more comfortable for Neolithic crops. ^15, 16^ Based on this, we divided our Central European group into Northern (above 50°N latitude) and Southern (below 50°N) populations. The Mesolithic hunter-gatherer population in Central Europe made a limited genetic contribution to the LBK population, whose members harbor only traces of hunter-gatherer admixture. ^7, 17, 18^ Contemporary populations from southeastern Europe have similarly low levels of hunter-gatherer ancestry. ^19^ In contrast, Neolithic southern European populations associated with the Cardial and Impressed Ware cultures followed a separate migration route (Figure 1), occupied a milder climate zone, and carried more Mesolithic ancestry. ^17, 20^ Individuals in this region tend to be shorter than those from Central Europe and combined with their admixed ancestry this has led to suggestions of a genetic basis for decreased statures in this region. ^7, 21^

**Figure 1:**
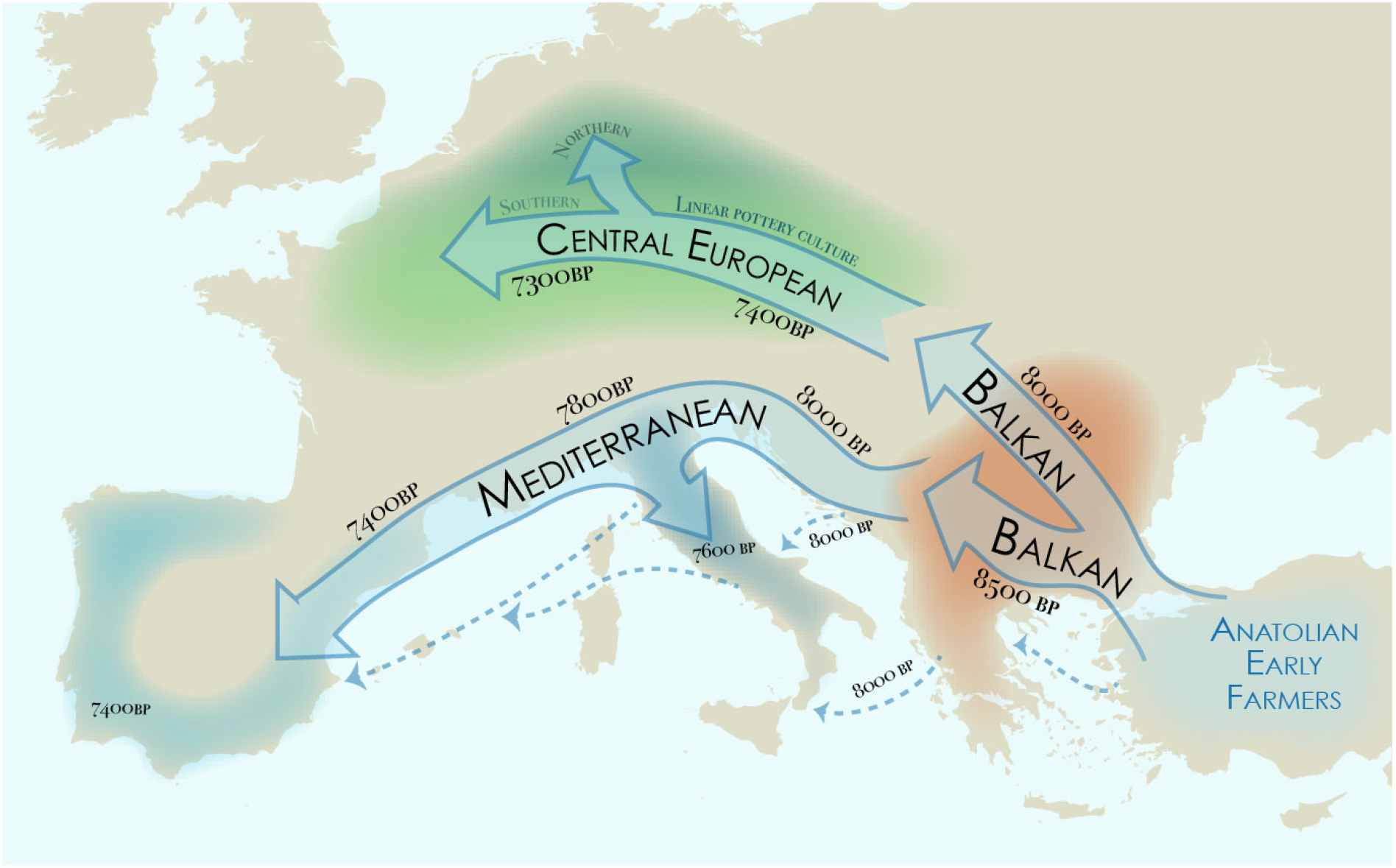
This schematic illustration highlights the two main migration routes from Anatolia to Europe during the Early Neolithic period. ^22^ Populations followed two routes: southern, along the Mediterranean coast (including sea routes, generalized here by dashed blue lines) where they admixed with existing hunter- gatherer populations; or northern, through the Balkans and into Central Europe, with only limited hunter- gatherer admixture. We analyze patterns within the Linearbandkeramik culture, dividing it into Northern and Southern Central European groups.

By comparing and contrasting these four closely related archaeological populations, we aim to investigate how differences in environment and genetics combine to produce observed phenotypes. We collected genetic data, skeletal metrics, paleopathology, and dietary stable isotopes to begin separating the effects of each on Neolithic stature trends. By specifically investigating and controlling for the effects of genetics in these samples, we are able to provide nuanced interpretations of height variation, gain a better understanding of the aspects of height which are controlled by genetics or environment, and show evidence for sex-specific cultural effects which modify the genetically predicted patterns. We illustrate the strengths of leveraging multidisciplinary datasets, and indicate caution when analyzing genotype-phenotype relationships without complete data, especially for traits which are not preserved in the archaeological record and cannot be directly tested. This integrated analysis highlights the role of plasticity in morphology, and establishes culturally mediated disparities at least as early as the European Neolithic.

## 2 Results

### 2.1 Distribution of stature, polygenic scores, and stable isotope values

We collected either genetic, dietary stable isotope, paleopathology, or skeletal metric data from 1282 individuals associated with the archaeological LBK culture in Northern and Southern Central Europe dated to between 7700-6900 years before present (BP), as well as 135 individuals from the southeastern (Balkan) and 160 individuals from the southern (Mediterranean) regions dated between 8000-6000 BP. All individuals included in the analysis have at least one of the four data types available (Materials and Methods; Figure 2; Supplementary Figure 1).

**Figure 2:**
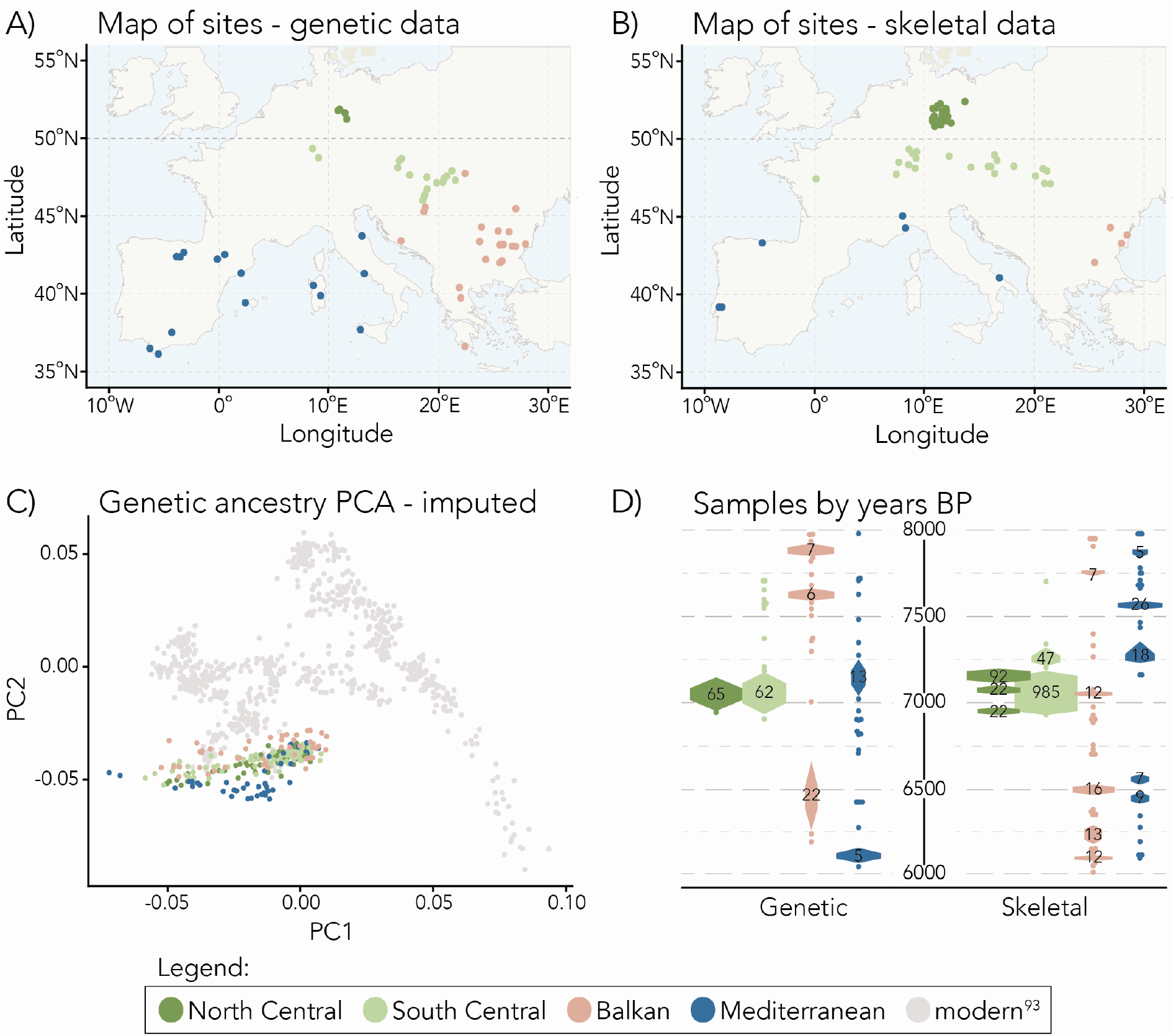
Upper row: Sites used for genetic (A) and skeletal (B) data in the analysis. The Central European population is split into Northern and Southern groups at 50°N latitude (emphasized). Lower row: (C) imputed genetic data projected into the PCA space of 777 modern Eurasian individuals (grey points). (D) sample numbers by years before present (years BP) for skeletal (right) and genetic (left) data.

Observed patterns of femur length vary between sexes and populations. Male femora show no apparent difference between the Central and Balkan regions (p=0.56), but Mediterranean males are significantly shorter (p=5.5*×*10*^−^*^7^, *β*=-1.44cm). Conversely, female femora show a different pattern, with no significant difference between Mediterranean, South Central (p=0.97), and Balkan (p=0.54) populations, but substantially shorter values in the North Central population (p=9 *×* 10*^−^*^0^^7^, *β*=- 2.0cm) (Figure 3A). Differences between male and female femur lengths are highly significant in all popultions (p<2*×*10*^−^*^1^^6^). In contrast to the differences in femoral lengths, polygenic scores for height are very similar between all populations (pairwise t-tests p > 0.9) using the clumping/thresholding PRS construction (Figure 3B). PRS constructed with LDpred show Mediterranean individuals to be shorter than the other populations (p=0.002; Supplementary Figure 5). However, PRS constructed using summary statistics derived from between-sibling analysis finds similar genetic values in all populations with both PRS construction methods, so we conclude that apparent lower Mediterranean PRS may be due to population stratification in the GWAS data and may not reflect a true genetic difference. There are no significant differences between male and female PRS in any population (Figure 3B), providing no evidence for a genetic basis to this dimorphism.

**Figure 3:**
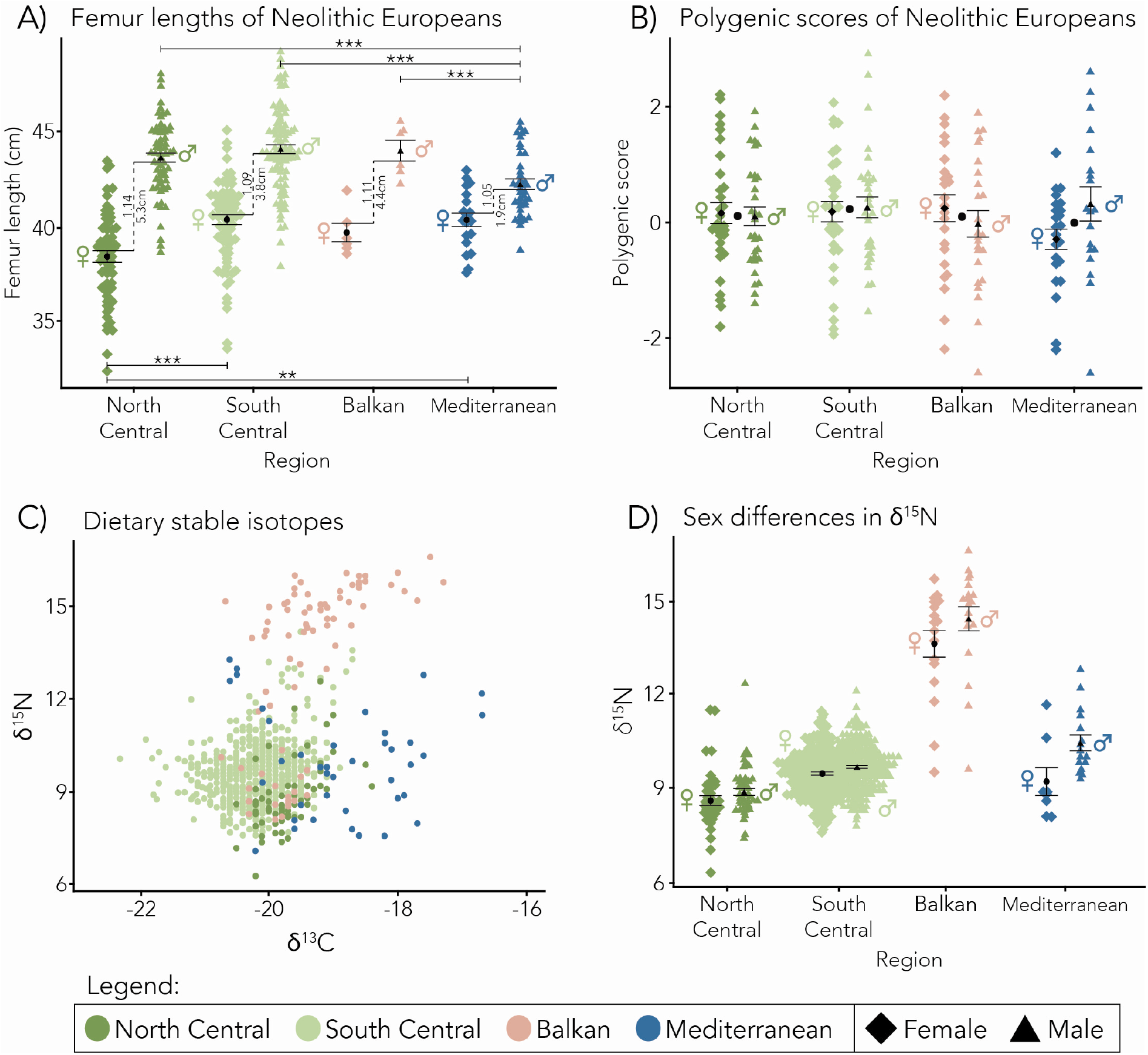
Solid bars across the tops of plots indicate significant differences between male populations by pairwise t-test; solid bars below plots indicate significant differences between female populations by pairwise t-test; p-values < 0.05 (*), < 0.01 (**) and < 0.001 (***); black points indicate the mean of each group; and vertical bars show mean standard error. A) Femur length in the four population: values to the right of the vertical dotted line are the difference between the mean male and female femora; values to the left are the sexual dimorphism ratios of male/female femur lengths for each population. B) Polygenic scores for the four populations show similar scores for individuals across all regions. Differences between male and female PRS are not significant. C) Plot of δ^13^C (x-axis) and δ^15^N (y-axis) dietary stable isotopes for the four populations: individuals from the Balkans are distinguished by high nitrogen values, while those in the Mediterranean generally have higher carbon. D) Sex differences in δ^15^N values by sex for each population: δ^15^N values are slightly higher for males in all populations, but this difference is only significant in the Mediterranean (p=0.035).

Signatures of *δ*^13^C and *δ*^15^N suggest different dietary patterns in each of the analyzed groups (Figure 3C, D). Both the Mediterranean and Balkan groups are significantly distinct from the Central in *δ*^13^C (multiple comparisons, maximum p=4.0 *×* 10*^−^*^1^^6^) and *δ*^15^N (multiple comparisons, maximum p=7.7 *×* 10*^−^*^1^^3^) values. Generally, the Balkan population is characterized by high *δ*^15^N values, while Mediterranean populations show high *δ*^13^C relative to the Central Europeans (Figure 3C). The exception to this pattern is a cluster of individuals, classified as Balkan in our analysis, which overlaps with the North Central population as well as some of the Mediterranean. These points represent individuals from present-day Greece and indicate that the diets of these peoples might better be classified as Mediterranean than Balkan. Nitrogen values are generally elevated in males compared to females (Figure 3D), but this difference is only significant in the Mediterranean (p=0.035).

### 2.2 Patterns of non-genetic factors in Central Europe

The most dramatic observation is the difference in female stature and consequent sexual dimporphism in Northern compared to Southern Central Europe. Female femora in the North are significantly shorter than female femora in the South (p=2.7 *×* 10*^−^*^6^, *β*=1.7cm), while male femora are highly similar (p=0.35) (Figure 3A). On average, male femora from the North are about 13% longer than female femora, Southern Central and Balkan male femora are about 9% and 11% longer respectively, and Mediterranean male femora are only 5% longer (Figure 3A). These values are reduced slightly when calculated using estimated statures instead of femora (North Central: 10%, South Central: 7%, Balkans: 8%, Mediterranean: 4%), possibly due to error associated with stature estimation (see Ref. 9) and body proportions, or because the relationship between femur length and stature is different between males and females. Where we have both genetic and metric data for the same individuals, there is a qualitative relationship between femur length and PRS; PRS tends to increase as femur lengths increase (Supplementary Figure 3B). However, the effect of PRS on femur length is only marginally significant (p=0.05), likely due to the small number of individuals with both types of data available in the sample (n=55).

Overall, trends in dietary stable isotopes show that both males and females in Southern Central Europe have significantly higher *δ*^15^N (male p=1.3 *×* 10*^−^*^9^, *β*=0.83‱; female p=5.3 *×* 10*^−^*^9^, *β*=0.87‱) and lower *δ*^13^C (male p=3 *×* 10*^−^*^4^, *β*=-0.30‱; female p=8.1 *×* 10*^−^*^7^, *β*=-0.38‱) as compared to the North. However, while males in both regions qualitatively have higher nitrogen, the interaction effect between sexes is not significant in either North or South (Figure 4A), indicating that the difference between male and female values in each region is not significant. There is no difference in carbon values between sexes. For individuals with both stature and stable isotope values, we find no statistically significant relationship between femur length and *δ*^15^N or *δ*^13^C in either Central group, separately or combined, but the sample is small.

**Figure 4:**
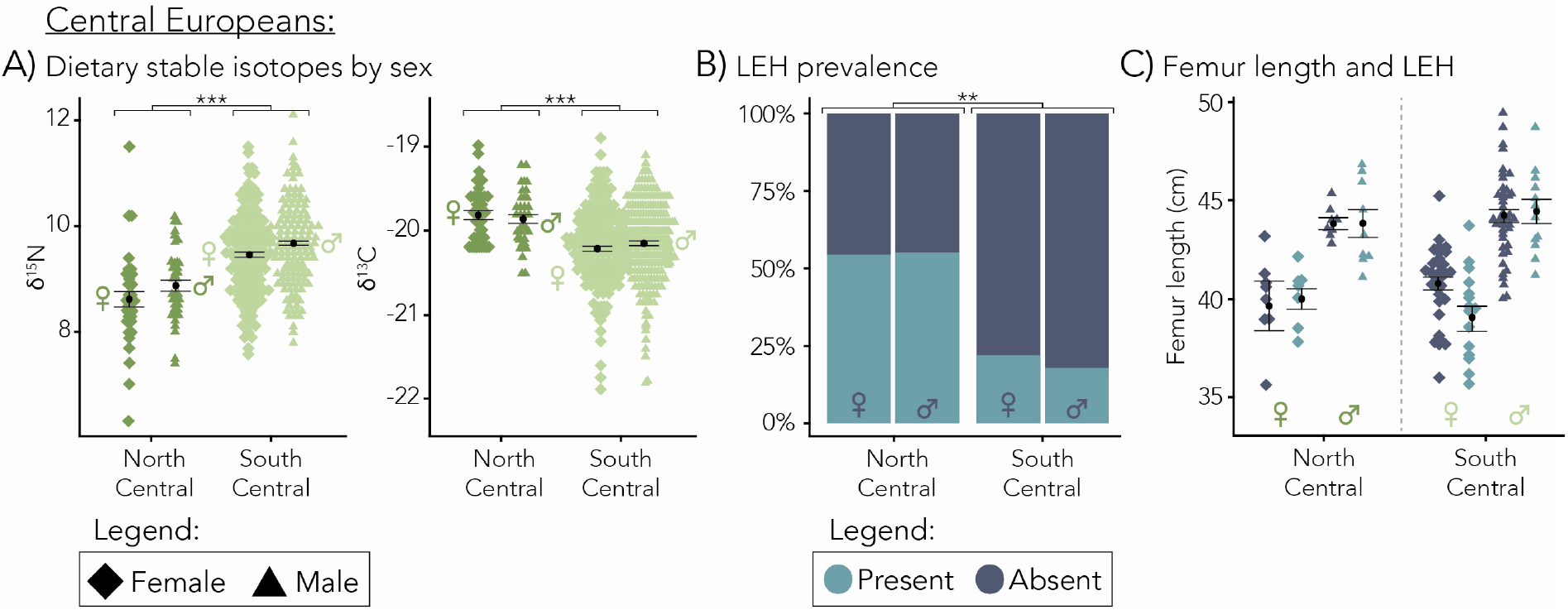
Evidence of environmental stress in Northern Central Europe. A) Differences in δ^13^C (right) and δ^15^N (left) values. Overall, the South has higher nitrogen values than the North (p=6.8 × 10^−1^^3^), and lower carbon (p=5.3 × 10^−1^^5^); within each population, the difference in isotopes between sexes is not significant. B) Proportion of linear enamel hypoplasias. The South has significantly less than the North (p=0.001). C) Presence of linear enamel hypoplasia is significantly associated with shorter femora (p=0.02); differences in prevalence between sexes are not significant.

We do, however, find a statistically significant relationship between presence of linear enamel hypoplasias (LEH) and shorter femora, suggesting that LEH may reflect an underlying variable in childhood that also affects stature (p=0.021, *β*=-1.0cm)(Figure 4C). Both males and females from the North are more likely to have LEH than individuals living in the South (p=0.002). Indeed, over 50% of the Northern sample have LEH while they are only present in about 20% of the Southern (Figure 4B). There is no significant difference between the number of males and females with LEH in either region. Though the interaction effect between sex and LEH on femur length is not significant, qualitatively the effect of LEH on femur length appears greater in females than in males (Figure 4C). When the sexes are analyzed separately, females with LEH do have significantly shorter femora than those without (p=0.018, *β*=-1.46), which is not the case for males (p=0.479). We hypothesize that the relationship between LEH and femur length is driven by females, but we lack an adequate sample size to detect the interaction effect in the full model. Incidence of cribra orbitalia is also significantly higher in the Northern region than in the Southern (p=1.8 *×* 10*^−^*^6^), though there is no relationship with femur length. There are no significant trends related to the presence of porotic hyperostosis.

In summary, comparison of Northern and Southern Central Europe identifies no predicted genetic difference in stature, which is consistent with male but not female femur length. This suggests a non-genetic basis for reduced female stature. Stable isotope data and skeletal stress indicators suggest lower protein intake and more general stress in the North; however, males and females overall appear equally affected by these variables. Despite a similar number of hypoplasias in both sexes, shorter femora in females suggest that increased general stress, due to other unmeasured environmental or cultural factors, leads to a female-specific reduction in stature.

### 2.3 Patterns of genetic ancestry in the Mediterranean

In contrast to Northern Central Europe, Mediterranean Neolithic males are shorter than other groups, but females are not. PCA indicates that individuals from the Central regions and the Balkans share similar genetic ancestry while those from the Mediterranean are distinct (Figure 2C; unimputed PCA in Supplementary Figure 4A), a difference known to be due to higher levels of hunter-gatherer ancestry in the Mediterranean. ^17^ We therefore additionally compared our samples to Mesolithic individuals of Western Hunter-Gatherer (WHG) ancestry, as well as individuals from early Neolithic Anatolia. These two groups represent source populations for the two largest ancestry components in Europe at this time. ^7, 17^

On the PCA plots of these extended data, Neolithic Anatolians cluster with the Central and Balkan groups. While Mediterraneans are near the farmer cluster, they are shifted towards the WHG (Figure 5A; unimputed PCA in Supplementary Figure 4B). ADMIXTURE analysis of all six populations supports this conclusion, showing significantly increased proportions of WHG ancestry in the Neolithic Mediterranean as compared with the other groups (maximum p=0.002 vs the Balkans, Fig. 5C). The average proportion of WHG ancestry in the Mediterranean is 11.4%; in the Balkans, 5.3%; in the South Central, 4.1%; and in the North Central, 1.1%. If there are significant PRS differences between Mediterranean and other populations, they are likely linked to this greater WHG ancestry and reflect genetic differences between WHG and other populations.

**Figure 5:**
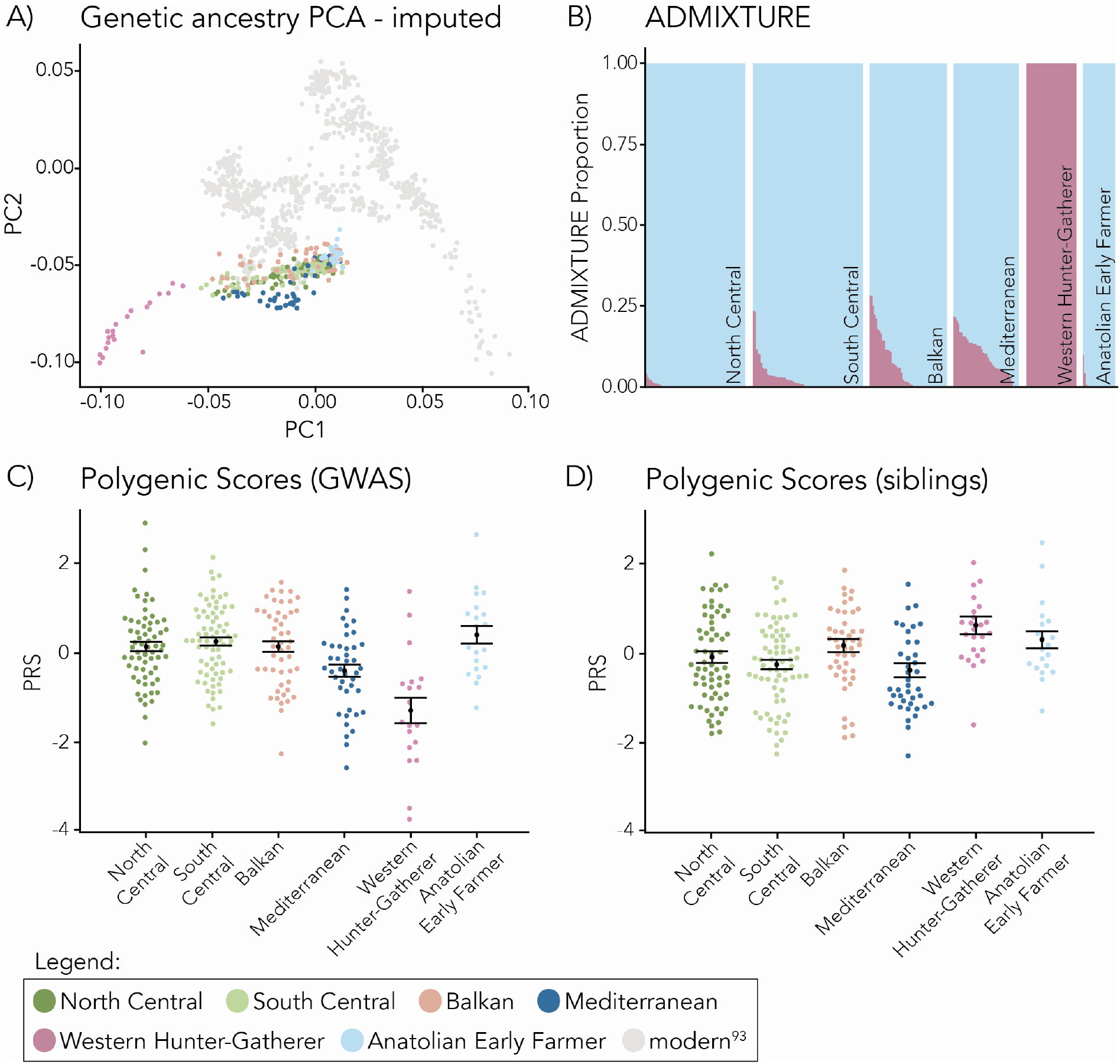
A) Ancient individuals projected into modern PCA space, including those of Mesolithic Western Hunter-Gatherer (WHG) and Anatolian Early Farmer ancestry. B) ADMIXTURE plot of K=2 ancestry groups showing the increased proportion of WHG ancestry in Mediterranean individuals. C) Polygenic scores for each region including Mesolithic Hunter-Gatherers and Anatolian Early Farmers. D) Polygenic scores calculated from between-sibling summary statistics.

Computing PRS using clumping/thresholding, we find that the WHG have the lowest PRS of any population in our data (maximum p=0.002 vs Mediterranean), while Anatolians are similar to the Balkan and Central Europeans. Among individuals, proportions of WHG ancestry are strongly associated with decreased PRS (p=1.6 *×* 10*^−^*^0^^6^, *β*=-0.08cm/%). However, when we compute PRS with an infinitesimal LDpred2 model, Mediterranean PRS is intermediate between Neolithic Europeans and WHG. When we repeat the LDpred analysis using summary statistics computed from between-sibling GWAS, ^23^ we find that the direction of Hunter-Gatherer PRS flips, and they have significantly greater PRS than the other groups (p=0.002) (Supplementary Figure 3A). The inconsistency of these results shows that the apparent PRS difference between WHG and Neolithic populations is highly sensitive to the PRS construction and summary statistics. This may indicate uncorrected population stratification in the non-sibling GWAS.^24, 25^ We therefore conclude that there is no strong evidence for a genetic difference in stature between Mediterranean and other Neolithic populations.

## 3 Discussion

Understanding the causes of past stature variation not only allows us to understand ancient communities, but may also provide us with insights into the origin and evolution of modern health patterns. However, interpretations of human stature variation through time remain confounded by the difficulty of separating genetic and environmental effects, obscuring trends. Recently, several researchers have begun to compile multivariate datasets for the purposes of understanding human stature. ^10, 26–28;^ however, many of these analyses do not directly take genetic effects into account, ^26, 28^ or cover very broad temporal or geographic ranges. ^10, 27^ In contrast, we aim to understand these processes on a finer scale to better interpret the outcomes of biological and environmental/cultural interactions. For instance, previous studies of stature variation from the Mesolithic to Neolithic indicated that Neolithic individuals were not achieving their genetic height potential, ^10^ but our analyses suggest that this effect might be heterogeneous, pertaining more to some locations and portions of society than others. Finally, while interesting in its own right, height can also serve as a model trait for how to incorporate genetics and anthropological data into studies of human morphology and variation. Here, by integrating genetic, cultural, and environmental data, we are able to begin teasing apart the contributions of genetic and non-genetic factors in producing the observed phenotypic variation. We also illustrate the existing limitations of interpreting genetic data.

Overall, the Central and Balkan groups are genetically homogenous with similar levels of huntergatherer admixture and polygenic scores, while Mediterranean individuals have more hunter-gatherer ancestry (consistent with previous observations ^7, 17, 18^). This may be associated with lower PRS, though this relationship is not robust and may simply reflect residual population stratification in the GWAS. None of our populations show evidence for substantial genetic differences in height between sexes (Supplemetary Figure 2), which is expected given that there is little evidence for sex differences in ancestry, or of sex-specific genetic effects on stature. ^29–31^ We can therefore largely exclude a genetic contribution to differences in stature between North Central individuals and other groups, while we find no strong evidence for a genetic contribution to shorter Mediterranean stature.

### 3.1 Sexual dimorphism in Central Europe reflects the effects of culture

Dietary differences between Southern and Northern Central European populations may indicate environmental stress in the North. In the early European Neolithic, the expansion of agriculture is thought to have been largely limited by poor soils and climate, as colder temperatures and decreased daylight made it increasingly difficult to grow early cereals (wheat, barley) and pulses (peas). ^32, 33^ The boundary to which these plants could be grown has been estimated to coincide with the northern limits of the LBK culture, ^13, 34^ and the majority of our Northern sites are concentrated near this climate edge in areas of fertile loess soils. ^14^ However, as there are many nuances which affect the interpretation of stable isotope values, especially between populations, differences between our Northern and Southern groups may not be as dramatic as they appear. An examination of isotope values from herbivorous animals in our study regions (using data from Refs. 16,35–39) indicates that baseline values of *δ*^15^N are elevated in the South Central region as compared to the North, potentially due to differences in climate and the use of manure as fertilizer. Therefore some portion of the difference between Northern and Southern nitrogen values might be attributable to variation in climate and farming practices rather than diet. Differences in carbon values between populations can be similarly sensitive to environment and we feel that interpreting the carbon results would be difficult without a more in-depth isotopic analysis. However, isotopic values from other studies show a higher proportion of plant vs animal foods in the North, particularly domesticated cereal grains. ^15, 36, 40^ Additionally, the available archaeological evidence supports some level of dietary difference between the Northern and Southern regions. While remains of cattle and dairy production are documented in Northern LBK sites, ^41, 42^ there is less archaeological evidence for the presence of other wild or domesticated animals that are seen in the South, indicating the people of this region were highly reliant on plant foods. ^40, 43^ We conclude that our observed differences in Northern and Southern stable isotope values probably reflect both dietary factors and differences in climate or farming practices.

It is therefore not surprising that people of the Northern Central region exhibit evidence of increased stress potentially due to unreliable and lower quality food resources. Lower protein consumption could be an indicator of dietary stress and has been linked to decreased stature.^44^ Diet can affect dimorphism in some cases, ^45^ but the isotopic signatures of males and females in the North Central, South Central, and Balkan regions are very similar, suggesting that this factor alone does not explain reduced female stature in the North. Femur length and isotope values for individuals are not significantly associated in our data, an indication that either diet has little effect on Neolithic stature or stable isotopes do not capture the elements of diet relevant to height. Alternatively, it is possible the range of variation in our data is too small to see this effect, or our sample of individuals with both metric and isotopic data is not large enough. In addition, we only analyzed adult samples and while the isotopic values of weaned children in the LBK fall within the range of adults, ^43^ it is possible that there could be sex differences in childhood diets affecting femur growth. Future studies incorporating collagen from long bones or teeth, rather than from ribs as we have here, would give dietary evidence with greater time depth, and might be able to provide more nuanced interpretations in the absence of a known-sex sub-adult population.

Paleopathological analysis also indicates increased stress in the Northern population in the form of increased incidence of linear enamel hypoplasia and cribra orbitalia. The causes of LEH formation are varied and their appearance in the bioarchaeological record is generally interpreted as a nonspecific indication of childhood stress. ^46^ Other archaeological sites have reported a high instance of LEH with high sexual dimorphism ratios in areas of Neolithic Europe, though the cause and meaning of these patterns was not explored (e.g. Ref. 47 and references therein). It has been suggested that cribra orbitalia might also reflect childhood stress, specifically anaemias, even when seen in adults. ^48^

Our results are consistent with others who have considered the same paleopathologies and found a qualitative relationship between presence of paleopathology and shorter femora. ^10^ In our data, incidence of both LEH and cribra orbitalia are higher in Northern compared to Southern Central Europe, but are not different between sexes in either group. The association between shorter femora and presence of LEH appears to be driven by females, suggesting a moderating factor causing a female-specific effect despite equal incidence of LEH in both sexes.

While we see a general increase in stress shared between sexes in North Central Europe, typical population-level stress responses usually show male vulnerability and female buffering effects. ^49–51^ Though the exact causes and mechanisms are not well understood, female biology tends to have a less extreme response, or is “buffered”, to many diseases ^52–54^ and environmental changes ^55^ compared to males. Our data indicate an opposite pattern in Central Europe, and no evidence of a variable which acts upon females alone. However, the Northern population shows sexual dimorphism that is extreme by present-day standards. In most modern global populations the ratio of male to female height is 1.06-1.08^56^ (ratios in Ref. 56 range up to 1.12, but population locations or cultural affiliations are not given, see Ref. 45), though it is difficult to know how to compare height versus femur length ratios as the transformation from metrics to stature scales differently in males and females. Based on 147 European individuals from the past 100 years (using data from Ref. 57), we find that the height ratio is very similar to the ratio of femur length—typically within 1%. We therefore conclude that dimporphism ratios in Southern Central (1.09) and Balkan (1.11) Europeans are elevated and the ratio in the North Central region is exceptionally high (1.14). Few modern populations have height dimorphism ratios as high as 1.10, and those that we could find in the literature come from India ^58^ and the United Arab Emirates, ^59^ both countries known for their cultural preferences and biases for male children. ^60^

We therefore hypothesize that the effects of high environmental stress in the North were modulated by culture. Other researchers have noted specific situations in which culture buffers males against environmental effects and creates vulnerability in females: there is an association between decreased female stature and polygyny in cultures around the globe; ^61^ female height was more influenced by economic conditions during infancy and early childhood than males in lower-class 19th-century Europe; ^62^ sexual dimorphism ratios in modern Chile decreased after the institution of social and government programs to combat gender inequality; ^63^ and 20th-century female stature decreased in India during times of environmental stress due to sexually disproportionate investment of scarce resources. ^60^ In LBK sites, strontium isotope values show that females are more likely to be non-local compared males, suggesting patrilocality and potential differences in cultural treatment of females. ^14, 64^ In parallel to our evidence for higher biological variation in females, ongoing discussion about the relationship between biological sex and the formation of gendered identities in the LBK suggests more variation in the roles and identities of females compared to males. ^65^ We therefore suggest that culturally mediated differences led to sex-specific stress responses in Neolithic Central Europe *via* cultural practices which either directly decreased female stature or, more likely, supported catch-up growth preferentially in males. Though dimorphism ratios in the South Central and Balkan regions are not as extreme as in the North, they are elevated and also consistent with this pattern of male bias, but response is likely less exaggerated due to lower environmental stress conditions.

### 3.2 Mediterranean differences may have both genetic and environmental bases

In the Early Neolithic Mediterranean population we see decreased male stature and low dimorphism ratios (1.05) relative to other Neolithic populations. Mediterranean populations are genetically distinct from other Early Neolithic groups with a higher proportion of WHG ancestry. In some analyses, WHG ancestry proportion correlates with lower PRS for height. However, PRS in the Mediterranean and WHG populations are sensitive to PRS construction method likely due to residual population stratification in the GWAS. These inconsistent results mean that we can neither confirm nor exclude the possibility of a genetic contribution to differences in stature between the Mediterranean and other Early Neolithic populations, though on balance we find the likelihood for a substantial genetic contribution to be low. Even if it were not, the genetic effects alone could not explain the reduced dimorphism ratio, emphasizing the need to also consider cultural/environmental effects.

While the dimorphism ratio in the Mediterranean Neolithic is low, it is not outside the range of present-day populations. ^56^ In fact, while males are relatively short, the longest average female femur lengths of our data are in the Mediterranean. This reduction in dimorphism is commonly seen in populations where the sexes experience an equal stress burden: as males tend to be more sensitive, decreasing their height, females are biologically buffered and stature remains consistent. ^49–51, 66^ Although we do not have paleopathological stress data for the Mediterranean individuals in our sample, published values for other Neolithic Mediterranean populations are generally similar to those for South Central Europe, ^67–69^ with exceptions. ^70^ Dietary isotopes indicate that the Mediterranean diet differs in some aspects, with increased *δ*^13^C values compared to the other Neolithic populations, but similar *δ*^15^N values. Our data indicates similar protein intake and low-level stress as other Neolithic populations, but do not suggest any clear hypothesis for the difference in male stature between the Mediterranean and other Neolithic groups. Possible differences in Mediterranean body proportions which are not captured by femur length should be mentioned as a caveat, though this likely would not be enough to account for the differences in stature compared to the rest of Europe, and would not affect observed dimorphism within the population. Our hypothesis is that the Mediterranean experienced similar levels of environmental stress as other Neolithic groups, but that they did not share the cultural practices which preferentially supported males and increased female vulnerability.

### 3.3 Conclusion

By integrating genetic and anthropological data, we are able to begin to understand the contributions of genetics and environment to human variation, allowing us to better interpret the genetic, environmental, and cultural landscapes of Neolithic Europe. Our results are consistent with a model in which sexually dimorphic differences in femur length are culturally and environmentally driven: relatively low dimorphism in the Mediterranean caused by female buffering to environmental stress and less cultural male preference, and high dimorphism in Northern Central Europe caused by the interaction of relatively high environmental stress and strong cultural male preference. Some analyses suggest that differences in average femur length between Central/Southeastern Europe and the Mediterranean are associated with differing genetic ancestries, but lack of robustness, uncertainty about the transferrability of polygenic scores, and questions of residual population stratification prevent us from interpreting this conclusively. In this study we focused on the European Early Neolithic because of its relative genetic, cultural, and environmental homogeneity, but, with more data, these methods could be extended to other populations, traits, and timescales to further explore the effects of human culture on biological variation. Using this approach, we gain a deeper understanding of the relationship between phenotypic plasticity, culture and genetic architecture, which constrain the mechanisms by which human biology adapts to environment.

## 4 Materials and Methods

We collected a combination of genetic, dietary stable isotope, skeletal metric, and paleopathological (stress) data from 1282 individuals from the Central European Early Neolithic associated with the archaeological LBK culture, approximately 7700-6900 BP (Figure 2, Supplementary Table 1). As there is archaeological evidence for broad regional variation within the LBK and our sampled sites form clear geographic groups, we divided these individuals into two regions based on geographical location, those to the north of 50°N latitude (North Central) and those to the south (South Central) (Figure 2A-B; North Central n=203, n femur length=133, n isotopes=100, n aDNA=67, n stress=83; South Central n=1067, n femur length=187, n isotopes=670, n aDNA=72, n stress=523). Each individual has at least one of the data types, and while some individuals have multiple data types, the overlaps are small (Supplementary Figure 1).

To provide wider context, we also compared Central individuals to other Neolithic populations from southern European (Mediterranean) and southeastern European (Balkan) regions, and restricted to individuals dated to 8000-6000 BP. We chose these regions as the Neolithic transition occurs at similar times and is associated with populations closely related to Central Europe. The acceptable date range for inclusion in the study was expanded from that which defines the LBK as these dates encompass comparable Early Neolithic phases in other parts of Europe while maximizing the number of eligible individuals. There could be a possibility that the later Balkan and Mediterranean individuals were more adapted to Neolithic life than the Central European groups, as these samples cover a longer time period, but we found no statistical within-population differences in our variables between the early and late ranges of our time span (minimum p=0.08). We excluded areas such as Scandinavia and Britain, where Neolithic technologies were not generally adopted until a later date. For the final analysis, we included 127 Mediterranean (n femur length=60, n isotopes=25, n aDNA=42) and 139 Balkan (n femur length=12, n isotopes=78, n aDNA=49) individuals (Figure 2). Unfortunately, there is a wide range of recording and reporting used for skeletal stress indicators, and it was not possible to build a statistically powerful dataset in these two populations for comparison; as a result, we did not analyze paleopathology in these populations. Finally, we collected genetic data from Mesolithic hunter-gatherer (n=25, 14000-7080BP, south of 48*^◦^*N) and Anatolian Neolithic (n=21) individuals for additional comparison.

### 4.1 Genetic data

We obtained genetic data for a total of 276 individuals. ^7, 18–20, 71–87^ Most data were generated by targeting a set of 1.24 million SNPs (the “1240k” capture reagent). ^17, 75^ For each individual, we randomly selected a single allele from each of the 1240k sites. Coverage in our dataset is low (median coverage=0.33; coverage above 0.60 n=71), and typically, it is not possible to directly infer diploid genotypes, potentially limiting PRS performance. Imputation of missing genotypes has been shown to help improve polygenic predictions for low coverage ancient samples, ^9^ and we therefore imputed diploid genotypes using the two-stage method described in that paper, restricting to SNPs in the 1240k set.

We calculated polygenic scores as previously described. ^9^ Briefly, we used standing height summary statistics generated by *fastGWA* from 456,000 individuals of European ancestry in the UK Biobank ^88^ for analyses of combined-sex PRS, and summary statistics from male- and female-only UK Biobank GWAS generated by the Neale Lab. ^89^ To test the potential effects of residual population structure in our data, we also computed PRS using additional summary statistics from a between-sibling GWAS (n=99,997). ^23^ We intersected the sites from each of these datasets with those on the 1240k array and then further restricted to HapMap3 SNPs (SNPs n=405,000). We computed polygenic scores using both a clumping/thresholding approach (*r*^2^=0.3, p-value cutoff=10*^−^*^6^, 100kb windows in *plink2* ^90^), and an infinitesimal *LDpred2* model using their pre-computed LD reference panel. ^91^ Finally, we computed polygenic scores using the --score command in *plink2*. In order to maximize the possibility of detecting sex-specific effects, we generated sex-specific PRS using three different approaches: 1) calculating PRS for all individuals using the female summary statistics; 2) calculating PRS for all individuals using the male summary statistics; and 3) calculating PRS for males and females separately using their respective summary statistics. While approach 3 seems at first to be the best for detecting these effects, observed patterns potentially become difficult to interpret due to differences in scaling between male and female PRS calculated as separate datasets. We computed principal components for both unimputed and imputed data using *smartpca*, ^92^ projecting ancient individuals onto principal component axes defined by 777 present-day West Eurasian individuals. ^93^ We also estimated K=2 unsupervised ADMIXTURE^94^ components for unimputed ancient individuals after first LD pruning using the command --pairwise-indep 200 25 0.4 in *plink2*.

### 4.2 Osteology and stable isotope data

We aggregated skeletal metric data from both published^57, 95–99^ and unpublished (n=28) sources. Maximum femur lengths were recorded when available, otherwise we estimated femur length from published stature estimates. ^9^ Estimated femur lengths correlate highly with stature estimates, but decrease the error that results from combining different estimations methods. The method from Ref. 100 provides separate equations for estimating the statures of northern vs. southern Europeans when using the tibia, due to differences in body proportions between the regions. There are two Mediterranean samples for which we estimated the length of the femur based on statures which used the southern tibia equation. Ref. 100 does not provide regional equations for femur estimation, so for these two individuals, we estimated femur length using the reverse of this region-agnostic femur equation.

For the individuals in this study who do not have genetic data, morphology was used to estimate sex. The majority of individuals have been taken from previous publications, and we used the sexes which had been estimated by those authors. For the individuals in our study which have not been previously published, sex was determined by co-authors using a 5-point scale on the cranium and pelvis as described by Ref. 101. For all individuals, sexes determined as probable male or probable female were coded in our study as either male or female as appropriate. Subadults and those with indeterminate morphologies were coded as NA, resulting in these individuals being dropped from the sex-specific analyses. The majority of sexes for individuals with metric data were determined by, or supervised by, co-authors and the remainder (n=13) either have genetic sexes or come from Ref. 57 which we consider a reliable source. Despite generally high accuracy for morphological sex determination, some level of uncertainty always remains, mainly due to variation in sexual dimorphism and preservation of the remains. ^102^ Sex estimations for our sample have all been performed in the last 20 years, and the majority within the last 5 years, meaning the researchers who performed them should be aware of avoiding the biases which can affect sex-ratios in the estimations of older data. Our dataset is large enough that small errors in classification of sex should not make substantial differences to results or interpretation, but the potential for inaccurate morphological sex estimations must always be considered in any osteological analysis. A large portion of our paleopathology data comes from tables S3 and S6 of Ref. 103, in which there are many instances of the same individual listed in both tables, but with discordant sex estimations. As we could not determine the reason for these discrepancies, we used the sex which was reported in the original publications cited as sources for their data. The few individuals (n=3) for whom this could not be resolved were treated as indeterminate and coded as NA. Ages were determined based on the average of the age range reported for each individual in their original publications.

For the paleopathological data in Central Europe, we took data from published sources, ^95, 96, 103–106^ as presence/absence of linear enamel hypoplasia (LEH), porotic hyperostosis, and cribra orbitalia. These three pathologies are often used by anthropologists as indicators of general, non-specific stress experienced by individuals or populations. While the exact eitiologies of these pathologies are generally not known, they have been shown to change through time within and between populations, and often correlate with environmental, social, or cultural shifts. Linear enamel hypoplasias are horizontal defects in tooth enamel that form during episodes of childhood stress severe enough to interrupt growth for some period of time, usually associated with dietary deficiency or infectious disease. ^46^ Individuals can exhibit one or multiple LEH on single or multiple teeth and in order to minimize errors from differences in reporting LEH in the literature, we have simply recorded whether an individual had any LEH (present) or none (absent). Porotic hyperostosis and cribra orbitalia are both porous lesions that are distinguished by their appearance on either the cranial vault or roof of the eye orbit respectively. The eitiologies of these are mostly unknown and though they are traditionally associated with amaemias, there are also a number of other conditions that can produce the same type of lesions. Medically, there is little evidence of these pathological changes despite their prevalence in the bioarchaeological record. ^107^ Similar to LEH, we have recorded these as either present or absent for each individual in order to standardize between reporting conventions across publications.

While sensitive to confounding factors such as climate, vegetation, and individual metabolism, ^108^ *δ*^13^C and *δ*^15^N stable isotope data can be used to reconstruct aspects of diet. ^109^ Here, carbon values are indicative of dietary plant resources and of the terrestrial vs marine vs limnic provenance of food, while nitrogen values are mainly associated with dietary protein intake and generally indicate proportions of plant- vs animal-based diets. ^108, 109^ We collected dietary stable isotopes *δ*^13^C and *δ*^15^N from published ^16, 35–37, 43, 95, 97, 103, 105, 106, 108, 110–113^ and unpublished (n=38) reports. We excluded atomic mass spectrometer (AMS) values derived from radiocarbon dating, as they may not be comparable to isotope-ratio mass spectrometer (IRMS) measurements, as well as values from children below the age of three, due to increased nitrogen values from breastfeeding. Stable isotope values from older children were included in population-wide diet analyses as the isotope ranges fall within those of adults; however, we only included adults with estimated sexes in the sex-based diet analyses. If information on the sampled material was available, we chose values measured from rib collagen, as these samples are most plentiful, though they only reflect the last few years of the individual’s life.

All previously unpublished osteological data was collected and analyzed by co-authors with permission from the necessary regulating organizations and in accordance with German laws and policies.

### 4.3 Statistical models

We tested the effects of PRS, femur length, and isotope data on stature using linear regression models including sex and geographic region as covariates in combination with other variables as appropriate (e.g., femur *∼* sex + region + PRS; *δ*^15^N *∼* sex + region + femur). We included interaction terms to test the relationships between geographic regions and sex (e.g., femur *∼* region * sex) and used t-tests to test within-sex differences between regions. We used logistic regression with the same covariates to test for factors affecting presence/absence of paleopathologies. We carried out all statistical tests using the base functions in R version 4.0. ^114^

### 4.4 Data Availability

All non-genetic data and polygenic scores used in this analysis are provided in Supplementary Table 1. Original ancient DNA data files can be downloaded from the resources provided in their cited publications. Previously published osteological data can be found in their cited sources (Supplementary Table 1).

### 4.5 Code Availability

R code used in this analysis is available at https://github.com/mathilab/Neolithic_height.git.

## Acknowledgements

This work was supported by a grant from the National Science Foundation [BCS2123627] to IM. Metric skeletal data collection from Stuttgart-Mühlhausen was funded by the Deutsche Forschungsgemeinschaft (DFG; German Research Foundation) [RO 4148/1-1]. The investigations of the skeletal collections from Derenburg, Karsdorf, and Halberstadt were supported by the DFG [Al 287/7-1 and 7-3; Me 3245/1-1 and 1-3]. The content is the responsibility of the authors and does not necessarily represent the official view of the NSF or other funders.

## 4.6 Author Contributions

S.L.C. designed the study, collected data, performed analysis, and wrote the manuscript; I.M. designed the study and wrote the manuscript; N.N. and K.W.A., contributed data and archaeological background; E.R. contributed data and performed analysis; M.F., J.W., H.M., and W.H. contributed data. All authors edited and approved the final version.

