## Supplemental figures for "Socio-cultural practices may have affected sexual dimorphism in stature in Early Neolithic Europe"

**Supplement**

Supp.Fig. 1: Data set counts

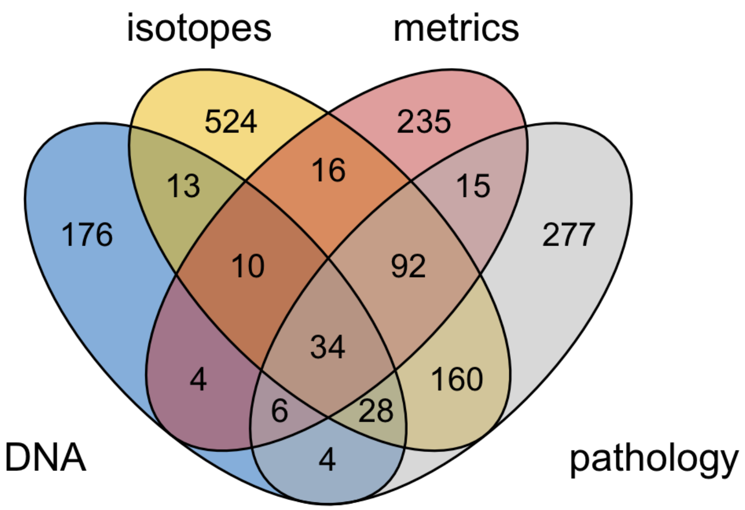


Figure S1: *Venn diagram illustrating the overlaps in data. All individuals have at least one type of data, but only subsets have more than one type.*

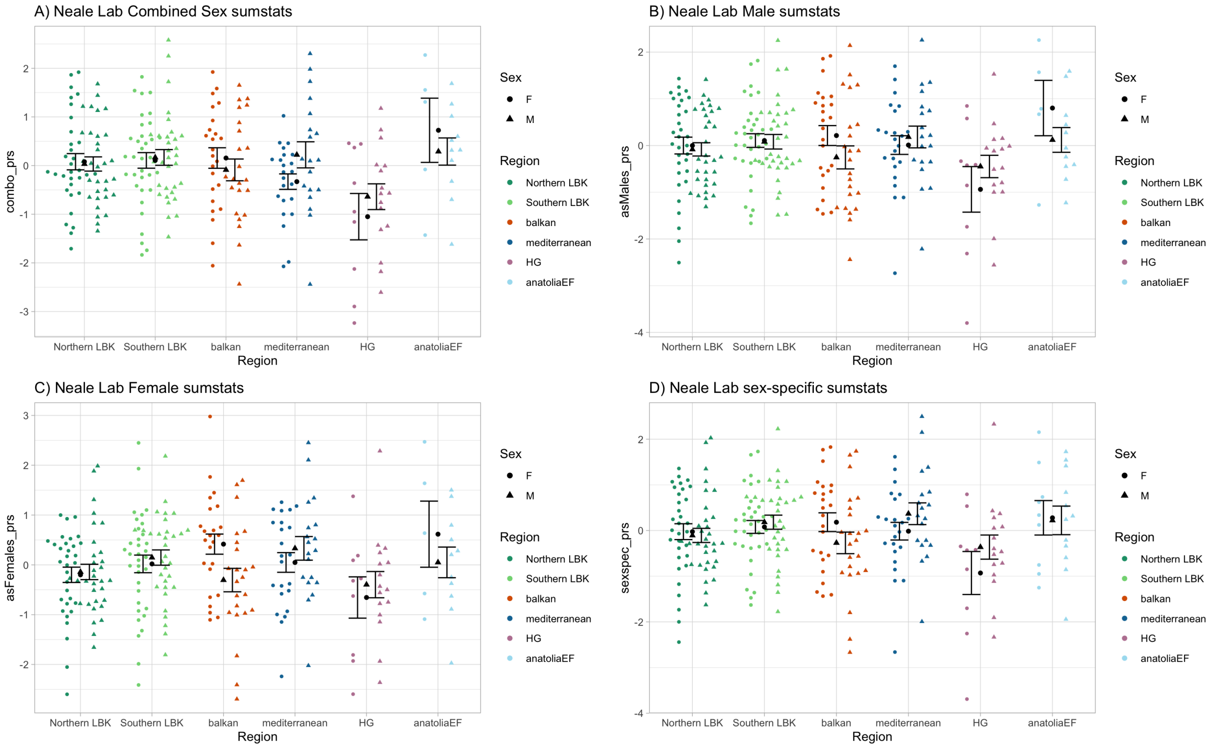


Figure S2: *Sex-specific clumping/thresholding PRS. A) PRS calculated using the combined sex summary statistics from the Neale Lab. There are no significant differences. B) PRS calculated for all individuals using the male-specific summary statistics from the Neale Lab. There are no significant differences. C) PRS calculated for all individuals using the female-specific summary statistics from the Neale Lab. There are no significant differences. D) PRS calculated for males and females separately using their respective summary statistics. There are no significant differences. (Summary stats: Neale Lab (2018))*

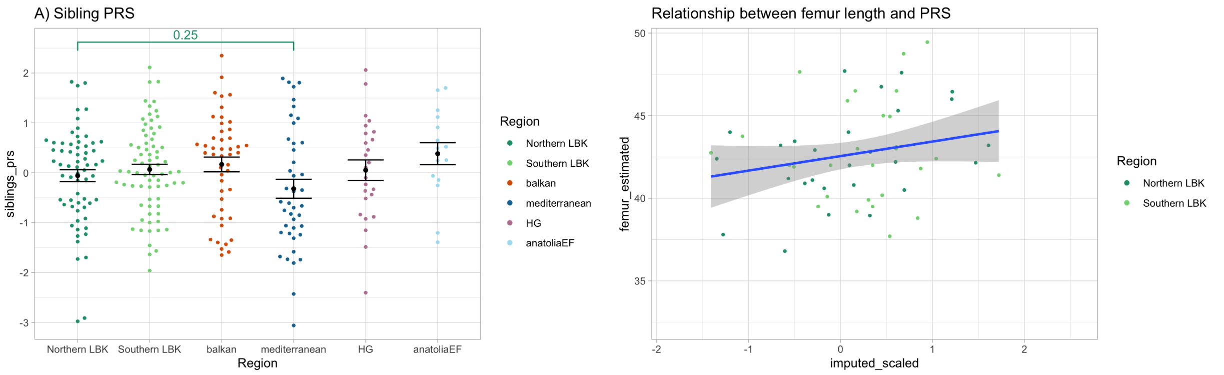


Figure S3: *A) PRS calculated with between-sibling effect sizes using the LDpred2 model. B) Femur length (y-axis) increases with clumping/thresholding PRS (x-axis) though the effect is not significant likely due to the small sample size.*

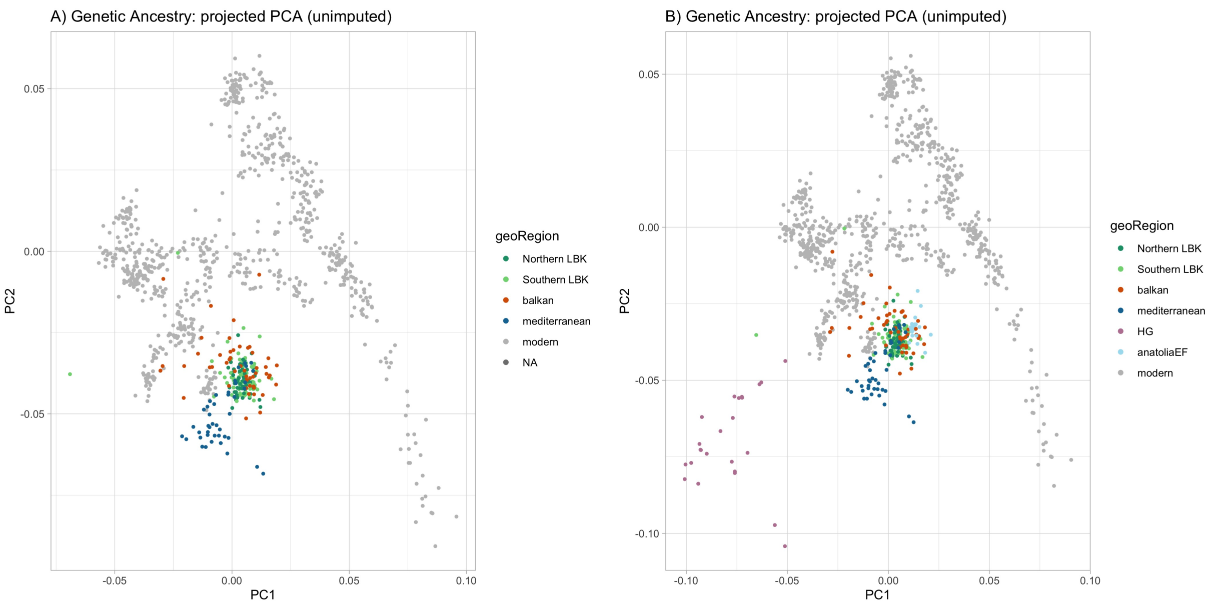


Figure S4: *PCA for unimputed DNA. A) Unimputed PCA for Neolithic Europe, equivalent to Figure 2C in the main text. B) Unimputed PCA for all samples, equivalent to Figure 5A in the main text.*

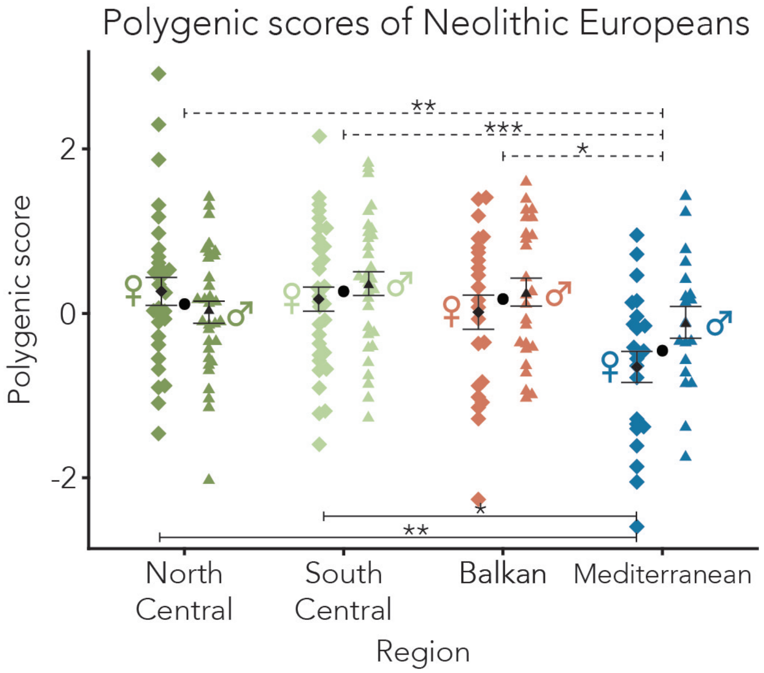


Figure S5: *LDpred2 PRS. Polygenic scores calculated using infinitessimal LDpred 2 model (Privé, 2020). There is no difference between North Central, South Central or Balkans populations. Mediterraneans PRS are significantly lower than the other groups (p=0.002). Similarly, the difference between males and females is only significant in the Mediterranean (p=0.04).*

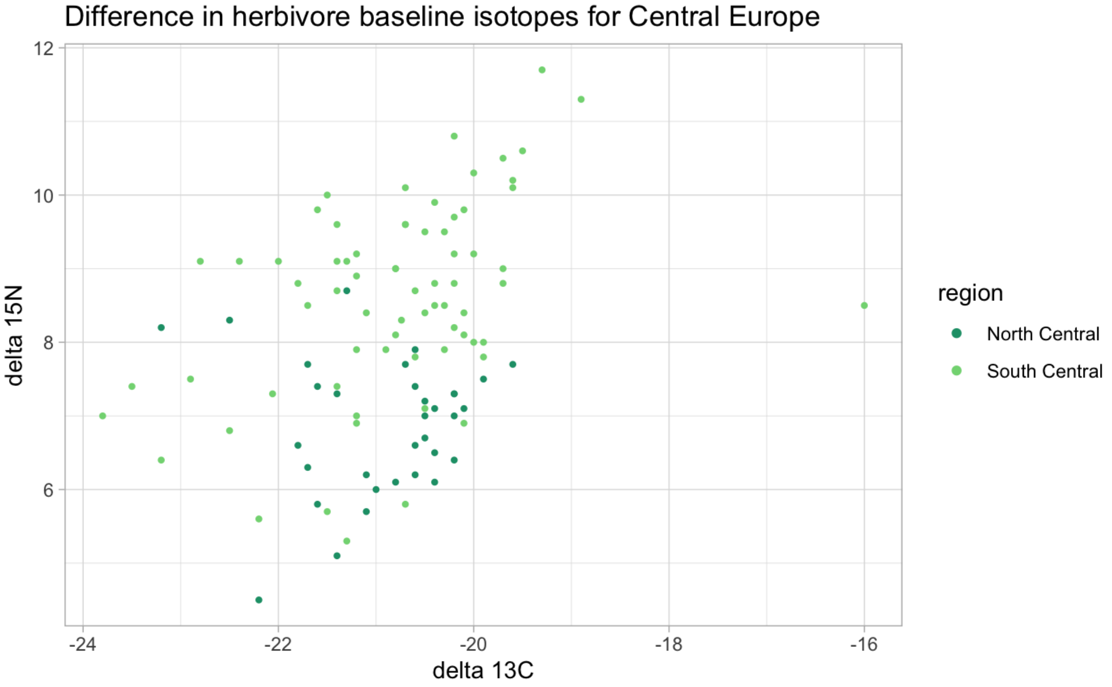


Figure S6: *Isotopic data from herbivores to establish a baseline for interpretation of human isotopic results. See citation in main text for data sources.*
